# Panton Valentine leucocidin enhances community-acquired methicillin-resistant *Staphylococcus aureus* colonisation of the gut

**DOI:** 10.1101/2020.07.10.157032

**Authors:** Florence Couzon, Nadège Bourgeois-Nicolaos, Yvonne Benito, Macarena Larroude, Anne Tristan, Jean-Philippe Rasigade, Binh An Diep, Florence Doucet-Populaire, François Vandenesch

## Abstract

**Objective:** Community-acquired methicillin resistant *Staphylococcus aureus* (CA-MRSA) independently emerged and became epidemic at the end of the 20^th^ century. Since gut carriage was reported for CA-MRSA and since the common feature of historical CA-MRSA is to harbour Panton Valentine leucocidin (PVL), the question of the possible involvement of this toxin in gut carriage was investigated in mice and cellular models.

**Methods:** CA-MRSA of three lineages (USA300, USA1100, and ST80) and their isogenic Δ*pvl* derivatives, were tested in competition for gut colonisation in mice and in a model of bacterial adhesion to mucus-producing intestinal epithelial cells.

**Results:** Mice inoculated with CA-MRSA and their Δ*pvl* derivatives had their gut successfully colonised by the three lineages regardless of the presence of PVL; however, the wild type (WT) CA-MRSA outcompeted the Δ*pvl* derivatives by at least 3 log after 40 days for all lineages tested. *In vitro* competition of CA-MRSA with their Δ*pvl* derivatives showed no fitness disequilibrium after 6 weeks, ensuring that the results obtained in mice did not result from direct bacterial interference. Direct fluorescence assay of mice intestine showed *S. aureus* localised at the mucosal surface of the intestine and within the intestinal crypts, but not within epithelial cells, suggesting a bacterial tropism for the mucus layer. Significant difference in adhesion to intestinal epithelial cells between WT and *pvl* knockout was only observed on mucus-producing cells, and not on non-producing ones.

**Conclusion:** PVL enhances CA-MRSA gut colonisation in mice by a mechanism involving adhesion-colonisation of the mucus layer.

## Introduction

*Staphylococcus aureus* remains one of the most common causative agents for both nosocomial and community-acquired infections. It colonises asymptomatically about one third of the human population and may cause mild to severe and occasionally life-threatening infections.^1^ Until the mid1990’s, methicillin-resistant *S. aureus* (MRSA) infections were reported exclusively from hospital settings and most hospital-associated MRSA (HA-MRSA) diseases resulted from a limited number of successful clones.^2^ However, in the beginning of the 21^st^ century, MRSA infections began to be reported in healthy individuals without risk factors or known connections to health care institutions.^3,4^ These community-acquired (CA)-MRSA strains had genetic backgrounds distinct from the traditional HA-MRSA strains with specific lineages predominating in different continents such as the ST8 SCC*mec*IVa pulsotype USA300 in the USA, the ST80 SCC*mec*IV in Europe, North Africa, and the Middle East, and the ST30 in Oceania.^5^ The genetic makeup of these CA-MRSA consists in the presence of a short SCC*mec*IV element as well as a prophage carrying the Panton Valentine leucocidin (PVL), an otherwise infrequent cytotoxin targeting the C5a receptor on human myeloid cells ^6^. A striking feature of these CA-MRSA clones was their emergence and spread in the community and their high epidemic success, notably that of the ST80 clone in North Africa,^7^ Greece,^8,9^ and the middle Est^10^ and that of the USA300 clone in the American continent.^11^ Knowledge on the underlying driving forces that rule the expansion of these clones is still limited; several non-exclusive contributing factors could be at play. An obvious contributing factor to MRSA dynamics is antibiotics use and overuse,^12^ and in this respect Gustave et al. showed that low-level antibiotics exposure in human and animal environments likely contributed to the demographic expansion of both European ST80 and USA300 lineages in the community setting.^13^ Moreover, the observation that PVL-positive CA-MRSA had apparently increased virulence in humans,^14^ notably for skin infections, suggests that higher bacterial load associated with increased severity of cutaneous infections could enhance dissemination between humans^13^. In addition to the aforementioned factors, the propensity to induce human carriage may be a determining role in the success of certain clones. Since intestinal and perineal carriage have been shown to contribute to environmental dissemination of *S. aureus*, patients with gut colonisation of *S. aureus* may serve as an important source of transmission.^15,16^ Indeed, extra-nasal carriage and multiple sites carriage (including pelvic sites and gut carriage) have been particularly described for CA-MRSA,^17–20^ suggesting that part of the epidemic success of these clones might be related to their ability to colonise extra-nasal sites. Given the fact that, besides the presence of SCC*mec*IV, the only common feature among the historical CA-MRSA (USA300, ST30, ST80) is the PVL-encoding bacteriophage,^4,5^ the possibility that PVL plays a determining role in gut colonisation is raised. The aim of the present study was thus to assess the propensity of CA-MRSA to colonise the gut in a mouse model and to evaluate whether the presence of PVL harboured by these CA-MRSA could play a determining role in gut colonisation.

## Material and methods

### Bacterial strains

CA-MRSA of three prevalent lineages (ST8-USA300, ST30-USA1100, and European-ST80) were selected and further referred to as wild type (WT) CA-MRSA. Their PVL knock-out derivatives (Δ*pvl*), obtained by allelic replacement, were either already available^21–23^ or constructed for the present study (Table 1). The BD0448 (USA1100 Δ*pvl*) strain was constructed from the BD0428 parental strain^23,24^ by transducing the *lukS/lukF*::*spc* mutation from the SF8300Δ*pvl* strain^25^ in which the *lukS*/*lukF* gene operon was replaced by a spectinomycin (*spc*) resistance gene using phage φ80α and selection on tryptic soy agar plate containing 1000 µg/mL spectinomycin. The successful transduction of *lukS*/*lukF*::*spc* mutation was confirmed by PCR and DNA sequencing using the previously described flanking primers X1 and X4.^26^

**Table 1.** Bacterial strains used in the study.

| Full Strain designation | Designation in article | Genetic background and modification | Relevant phenotype | Reference or source |
| --- | --- | --- | --- | --- |
| HT200020209/<br>LUG1799 | ST80 | ST80 MRSA-IV<br>European CA-MRSA | PVL+, OxaR, KanR | 21 |
| LUG1800 | ST80 $\Delta pvl$ | LUG1799 $\Delta pvl::tetR$ | PVL-, OxaR, KanaR, TetraR | 21 |
| LUG2216 | ST80-44 | LUG1799 after 44 days<br>in mice gut | PVL+, OxaR, KanR | This study |
| LUG2217 | ST80-44 $\Delta pvl$ | LUG1800 after 44 days<br>in mice gut | PVL-, OxaR, KanaR, TetraR | This study |
| ST20111292/<br>LUG2012 | USA300 | ST8 MRSA-IV USA300<br>North American variant<br>ACME+ | PVL+, OxaR, CipR | 22 |
| LUG2040 | USA300 $\Delta pvl$ | LUG2012 $\Delta pvl$ | PVL-, OxaR, CipR | 22 |
| LUG 3000 | USA300<br>$\Delta pvl-c$ | LUG2012 $\Delta pvl$, ORF29-30::*cat* | PVL-, OxaR, CipR,<br>CmR | This study |
| ST20101391/<br>LUG2282/BD428 | USA1100 | ST30 MRSA USA1100<br>Southwest Pacific Clone | PVL+, OxaR TetraR | 23 |
| ST20101393/<br>LUG2283/BD448 | USA1100<br>$\Delta pvl$ | LUG2282 $\Delta pvl::specR$ | PVL-, OxaR TetraR<br>SpecR | This study |

Insertion of the chloramphenicol resistance gene from pC194 plasmid^27^ in non-essential ORFs of USA300 Δ*pvl* (ORF SAUSA300_0087 to SAUSA300_0089 of USA300 FPR3757 homologous to Newman strain ORF NWMN29 and NWMN30),^28,29^ was achieved using the pMAD allelic replacement mutagenesis system^30^ and the primers shown in Table 2.

**Table 2.**
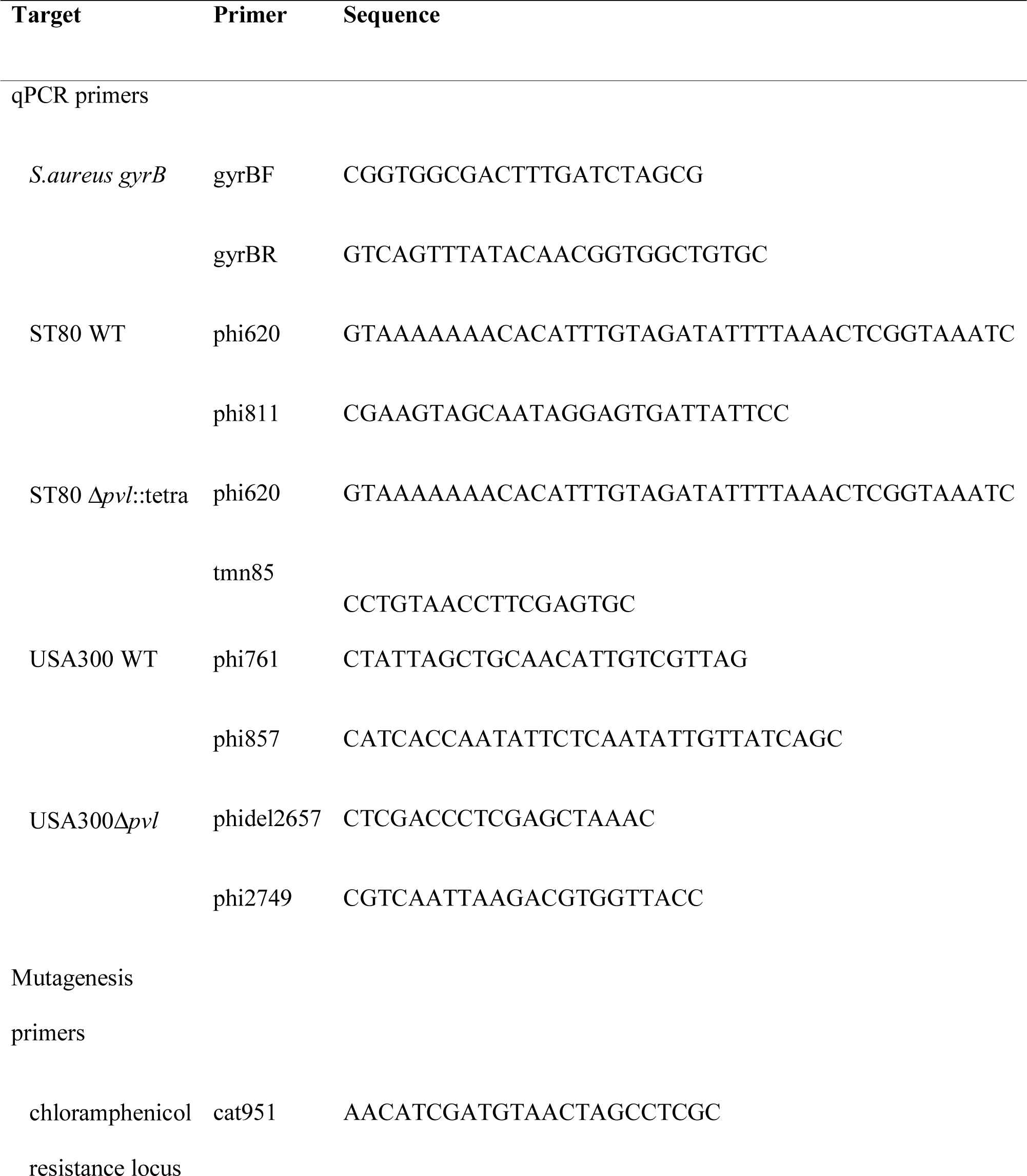
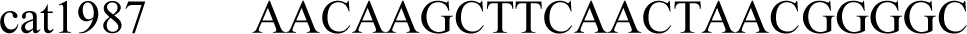
Primers used for the present study.

### Cells and Culture Conditions

The human intestinal epithelial cells HT29 ATCC HTB38^31^ were cultured in Dulbecco’s Modified Eagle’s Medium (DMEM, Thermofisher) supplemented with 10% foetal bovine serum (Gibco), at 37°C under 5% CO_2_ atmosphere. HT29-MTX10-6 (CelluloNet Biobank BB-0033-00072) cell population was derived from HT29 cells by adaptation to 10-6 Methotrexate^32^. HT29-MTX cells form a homogeneous population of polarised goblet cells that secrete mucins of gastric immunoreactivity. A mucus gel becomes visible on the cell surface of the cell layer at post-confluency, between 21 and 28 days of culture. These cells maintain their ability to differentiate under normal culture conditions (without MTX). They were cultured in DMEM supplemented with 10% foetal bovine serum and 1% sodium pyruvate 100mM (Thermofisher), at 37°C under 5% CO_2_ atmosphere.

### Axenic mice model of *S.aureus* gut colonisation

*In vivo* fitness was measured by co-colonisation experiments in a mouse model. To increase sensitivity, the latter was established in axenic female mice (C3H, *Cryopreservation, Distribution, Typage et Archivage Animal*, CNRS, Orléans, France). Mice (6-8 weeks old, 20 g) were housed in sterile isolators, as previously described.^33^ “Germ-free” mice underwent simultaneous intragastric inoculation of 10^8^ CFU of a mix in equal numbers of the 2 isogenic strains in a 500µl volume. Fresh faecal samples collected 3 times a week during 45 days were weighed, homogenised in PBS (0.1 mg/mL), and diluted 10 to 10. *S. aureus* was numbered by plating on Trypticase soy agar (TSA) with or without tetracycline at 0.4μg/ml for ST80, on TSA with or without spectinomycin at 1mg/ml for USA1100, and by plating on TSA without antibiotics followed by quantitative PCR (see below) for USA300. In the latter case, the respective CFU of WT and Δ*pvl* was inferred from the total CFU count on plate, distributed between WT and Δ*pvl* according to qPCR ratio. The remaining pellets were kept at −80°C for further analysis. At the end of the challenge, mice were sacrificed, the caecum was removed and immediately immersed in PFA 4% for at least 24hours.

### Relative DNA quantification

Mouse faeces were collected and frozen until DNA extraction. Total DNA was extracted using QuickGene DNA tissue kit S and the nucleic acid isolation system QuickGene Mini80 (Kurabo, Médiane diagnostics, Plaisir, France) according to the manufacturer’s recommendations. Prior to proteinase K digestion, each frozen faeces was crushed in lysis buffer in a 1.5 ml microcentrifuge tube using a pestle (USA scientific).

DNA was used as a template for the real-time PCR amplification using a LightCycler Nano instrument (Roche diagnostics), Fast Start Essential DNA Green Master Green (Roche diagnostics) and specific primers targeting *gyrB*, WT, and Δ*pvl* strains (Table 2). The relative amounts of WT/Δ*pvl* strains were determined by quantitative PCR and expressed as fold-change relative to the *gyr*B internal standard as described elsewhere.^34^ The relative DNA quantification was processed in duplicate for each of the faeces samples and corresponding isolates obtained.

### Direct fluorescence assay (DFA)

Intestinal tissues stored in PFA were then embedded in paraffin blocks and cut into 6 μm slices. Slides were deparaffinised, cleared in xylene, and rehydrated in preparation for staining, blocked with 1% bovine serum albumin (BSA) in tris-buffered saline (TBS) supplemented with 10% of monkey serum, for 2 hours at room temperature. Bacterial localisation was detected with primary anti-*S. aureus* (ab37644, Abcam, 1:100 dilution, overnight, at +4°C) then a secondary anti-mouse alexa fluor 488 labelled antibody (ab150101, Abcam, 1:1000 dilution, 1hour, at room temperature). Nuclear counterstain with DAPI was realised. Fluorescent images of gut intestine, using 488 nm (*S. aureus*) or 405 nm (DAPI) filters, were obtained under confocal microscopy Leica SP5 X (Leica, Solms, Germany), 63× objective under oil-immersion and analysed using Leica software LAS AF Lite (Leica Application Suite 2.6.0).

### *In vitro* competition

Equal numbers (10^3^ CFU in a 50µl volume) of WT and Δ*pvl* derivatives were inoculated in a tube containing 5ml-volume of Tryptic Soy Broth (TSB) and incubated at 37°C with shaking at 200 rpm during 24 hours. To facilitate the follow-up of the competition by using CFU counting on agar plates, a chloramphenicol resistance marker was introduced in USA300 Δ*pvl* whilst ST80 Δ*pvl* and USA1100 Δ*pvl* already harboured a tetracycline or a spectinomycin resistance marker, respectively (Table 1). Daily passage in fresh media was performed for 45 days by inoculating 10µl of the mixture to 5ml fresh TSB. *S. aureus* was numbered by CFU plating on TSA with or without tetracycline at 0.4 μg/ml for ST80, on TSA with or without chloramphenicol at 10µg/ml for USA300, and on TSA with or without spectinomycin at 1mg/ml for USA1100 to distinguish between isogenic strains.

### Adhesion to intestinal epithelial cells

HT29 cells were seeded (500,000 cells/ml) into 24-well culture plates and incubated in culture medium at 37°C in 5% CO_2_ for 24 hours until 80% confluence. HT29-MTX cells were seeded (100,000 cells/ml) into 24-well culture plates and incubated in culture medium at 37°C in 5% CO_2_ for 21 days until 100% confluence and mucus layer production. Bacterial cultures (WT and Δ*pvl*; 8 hours of growth) were washed with phosphate-buffered saline (PBS) and resuspended in fresh culture medium to achieve a multiplicity of infection of 10, subsequently confirmed by CFU counting upon agar plate inoculation. Cells were washed twice with 1 ml of PBS before the addition of bacteria. Cell cultures were then incubated at 37°C to allow for bacterial adhesion. After 2 hours, cells were washed twice with 1 ml of PBS and unbound bacteria were discarded. For the adhesion assay, infected cells were lysed using osmotic shock in pure sterile water. Viable adherent bacteria released after host cell lysis were enumerated by serial dilution and plate counting on agar plates.

### Statistical Analysis

The statistical analyses were performed using GraphPad Prism 8 software and R software v3.6.1. Data from 2 groups (WT and Δ*pvl* strains) were compared using Student two-tailed *t* test for paired samples. The significance threshold was set at 0.05. We used multivariate modelling to delineate the respective influences of *pvl*, mucus presence and genetic background on *S. aureus* adhesion in *in vitro* experiments. The % adhesion, transformed to log-odds, was modelled using mixed-effect linear regression adjusted for between-experiment variation using a random intercept per independent experiment. A first model was constructed using *pvl*, mucus and genetic background as predictor to test the independent effect of *pvl*. A second model also included an interaction term between *pvl* and mucus to test whether *pvl*-associated adhesion was dependent on the presence of mucus. Model coefficients are reported as odds-ratios of adhesion with 95% confidence intervals.

### Ethics statement

All mouse protocols were carried out in strict accordance with the Directive 2010/63/EU revising Directive 86/609/EEC on the protection of animals used for scientific purposes. This directive was translated in the French regulation as the Decret n°2013-118 of Feb 2013 under the jurisdiction of the Ministry of Education, Research and Technology.

The animal experiments were approved by the Committee on the Ethics of Animal Experiments n°26 of the University of Paris-Sud (agreement number 2012-120).

## Results

Both the CA-MRSA WT and Δ*pvl* derivatives of all three lineages tested showed initial success in colonising the mice gut. However, between two and three weeks after the start of the experiment, the Δ*pvl* strains declined whilst the WT CA-MRSA remained present in the gut at very high concentration (Figure1); after 6 weeks, there was a 3 log (USA1100 and USA300) to 5 log (ST80) decrease between WT and Δ*pvl* colonisation (Figures 1A-C).

**Figure 1.**
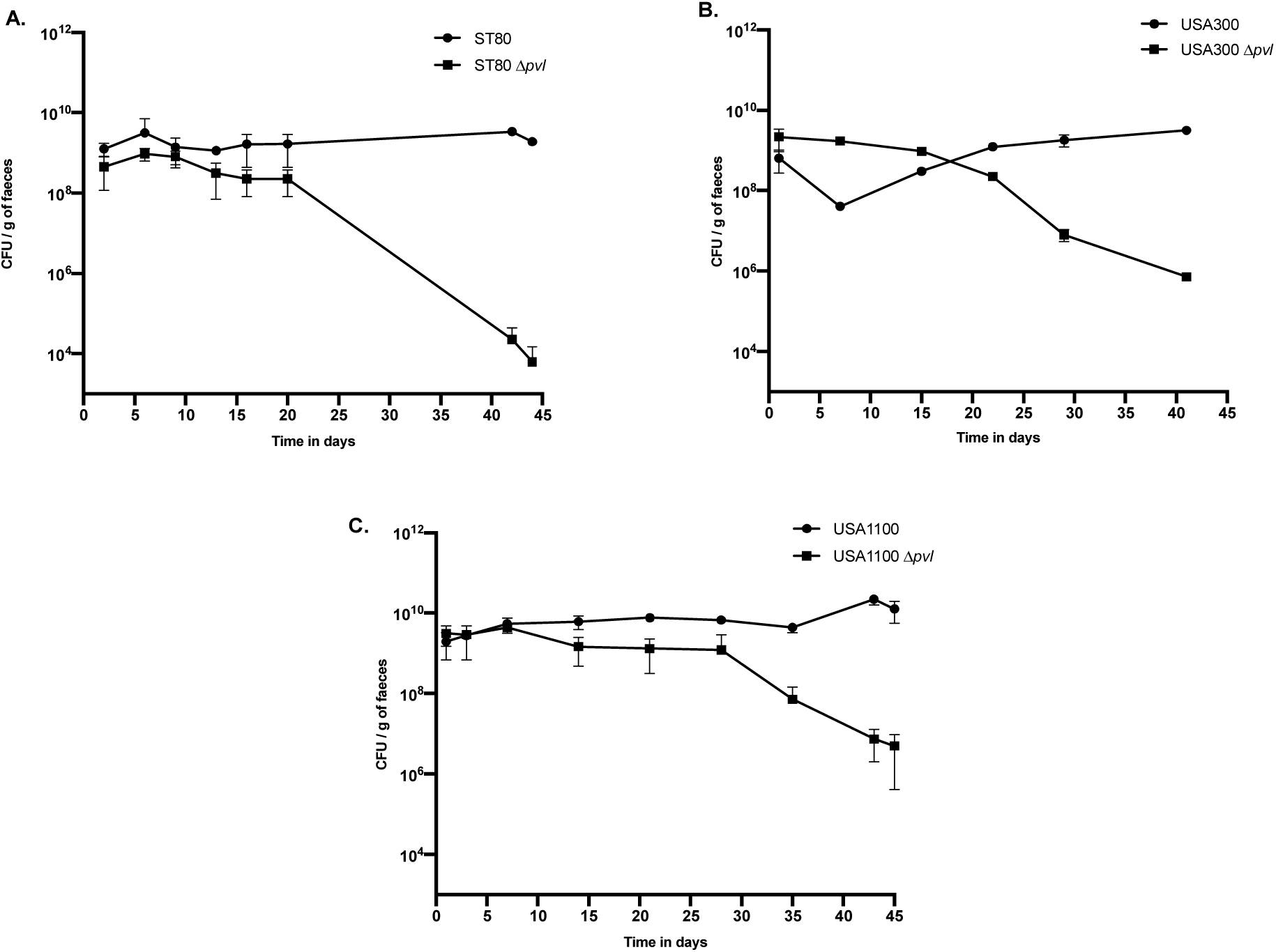
Axenic mice gut colonisation by CA-MRSA wild type and Δ*pvl* derivatives in dixenic challenges. ST80 WT and Δ*pvl* (Panel A; n=4), or USA300 WT and Δ*pvl* (Panel B; n=3) or USA1100 WT and Δ*pvl* (Panel C; n=4) were simultaneously intragastrically inoculated at day 0 to axenic mice using 10^8^ CFU of a mix in equal numbers of the two strains in a 500µl volume. The respective number of each strain in faeces was assessed by plate counting on antibiotic-containing media (ST80 and USA1100) or by plate counting without antibiotics and quantitative PCR (USA300). The results are expressed in CFU/g of faeces. In the case of USA300, the respective CFU of WT and Δ*pvl* was inferred from the total CFU count on plate, distributed between WT and Δ*pvl* according to qPCR ratio. Error bars show SEs.

Upon sacrifice of the mice after the challenge, DFA was performed to localise the fluorescent bacteria in the intestine of colonised mice. *S. aureus* was found on the surface of the intestinal epithelium in the mucus and also within the intestinal crypts (Figure 2). Of note, no fluorescent signal was observed within the cytoplasm of epithelial cells suggesting that colonisation of the gut by *S. aureus* did not reach an invasive step. Given the ratio of enumeration (Figure 1), the bacteria visible by DFA are very likely the WT (PVL positive).

**Figure 2.**
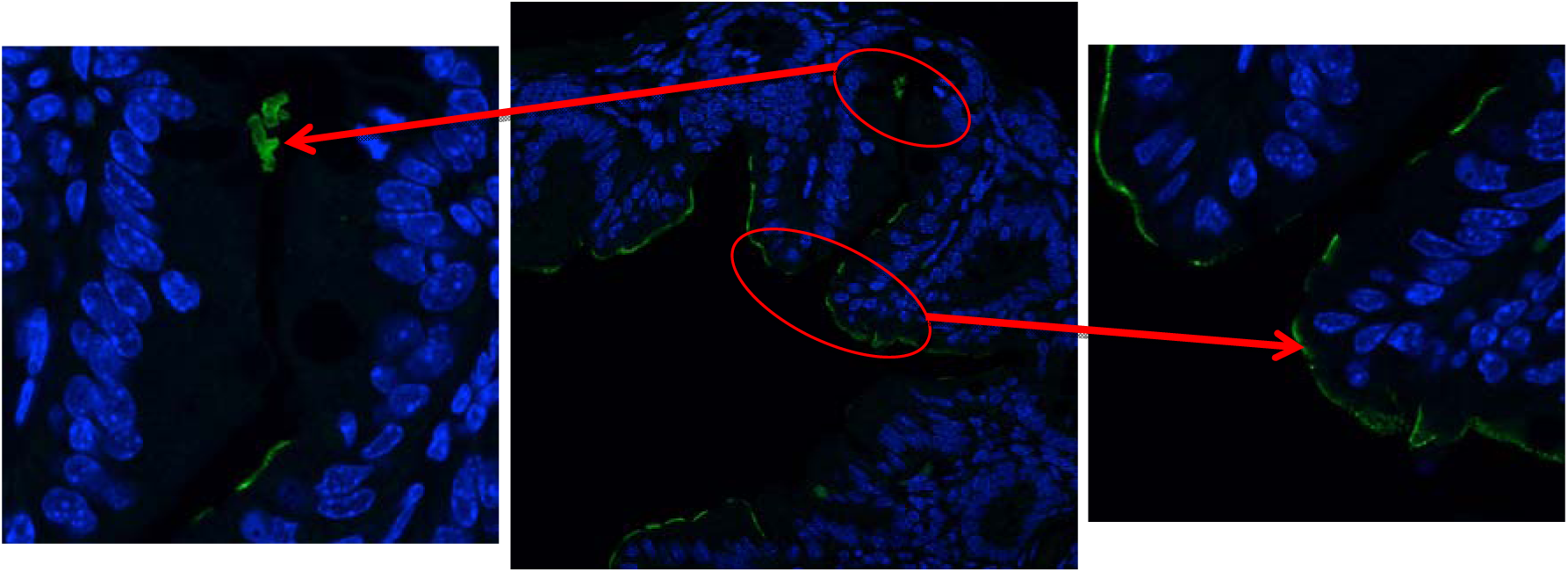
Direct fluorescence assay localisation of *S. aureus* on paraffin sections of mouse intestine. Paraffin sections prepared for histology were incubated with primary anti-*S. aureus* antibody followed by a secondary anti-mouse alexa fluor 488 (green) labelled antibody. Nuclear counterstain with DAPI (blue) was realised. Fluorescent images were obtained using confocal microscopy; regions of interest (circled) are enlarged and indicated by arrows.

To rule out the possibility that the Δ*pvl* strains were outcompeted because of a direct bacterial interference between WT and mutant strains, an in vitro competition assay was performed by inoculating the WT and Δ*pvl* strains at equivalent concentrations in BHI broth and by performing daily subcultures in fresh medium for 6 weeks. The results showed no fitness disequilibrium after 6 weeks for the three lineages (Figures 3A-C) confirming that the results obtained in mice did not result from direct bacterial interference. In addition, the same experiment was performed using strains ST80 WT and Δ*pvl* isolated after 6 weeks in mice (Figure 3D). Again, there was no fitness disequilibrium between strains after intestinal colonisation of mice.

**Figure 3.**
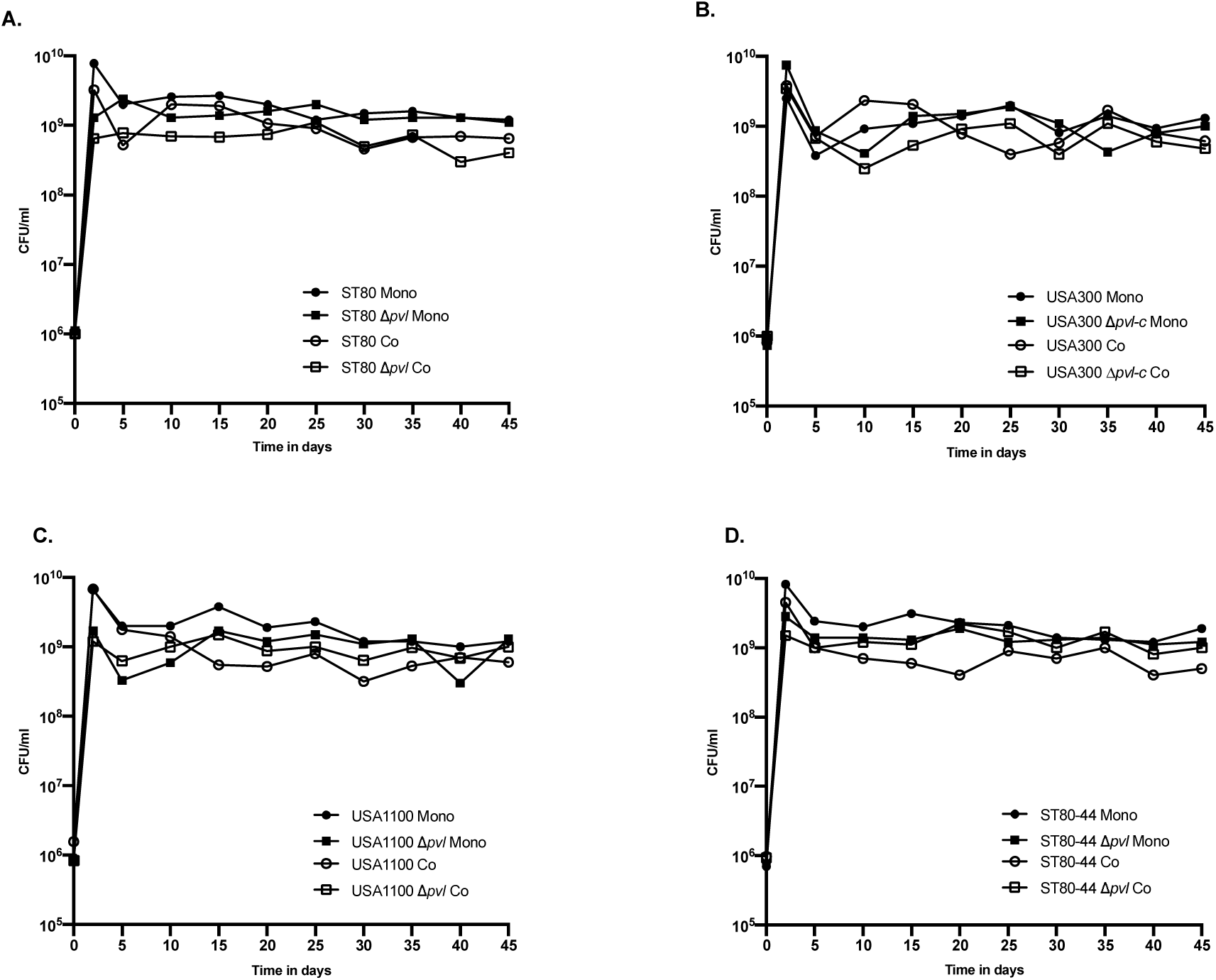
*In vitro* competition of CA-MRSA wild type and Δ*pvl* derivatives. Equal number (10^3^ CFU in a 50µl volume) of WT and Δ*pvl* derivatives of ST80 (panel A), USA300 (panel B), and USA1100 (panel C) in mono- and in co-culture were subcultured daily in TSB for 45 days. The same experiment was performed using strains ST80 WT and Δ*pvl* isolated after 6 weeks in mice (ST80-44 and ST80-44 Δ*pvl*, panel D). *S. aureus* was numbered by CFU plating on TSA with or without tetracycline for ST80, chloramphenicol for USA300, and spectinomycin for USA1100, to distinguish between isogenic strains. The results are expressed in CFU/ml.

In order to confirm the localisation of *S.aureus* on the surface of intestinal cells and to highlight a potential role of PVL in mucus adhesion, bacterial adhesion to mucus-producing intestinal epithelial cells was assessed for the three lineages and their Δ*pvl* derivatives. While there were no significant difference between WT and Δ*pvl* bacterial adhesion on non-mucus producing cells (HT29), adhesion on HT29 MTX was significantly higher for *S.aureus* WT, as compared to Δ*pvl*, for all three lineages tested (Figure 4).

**Figure 4.**
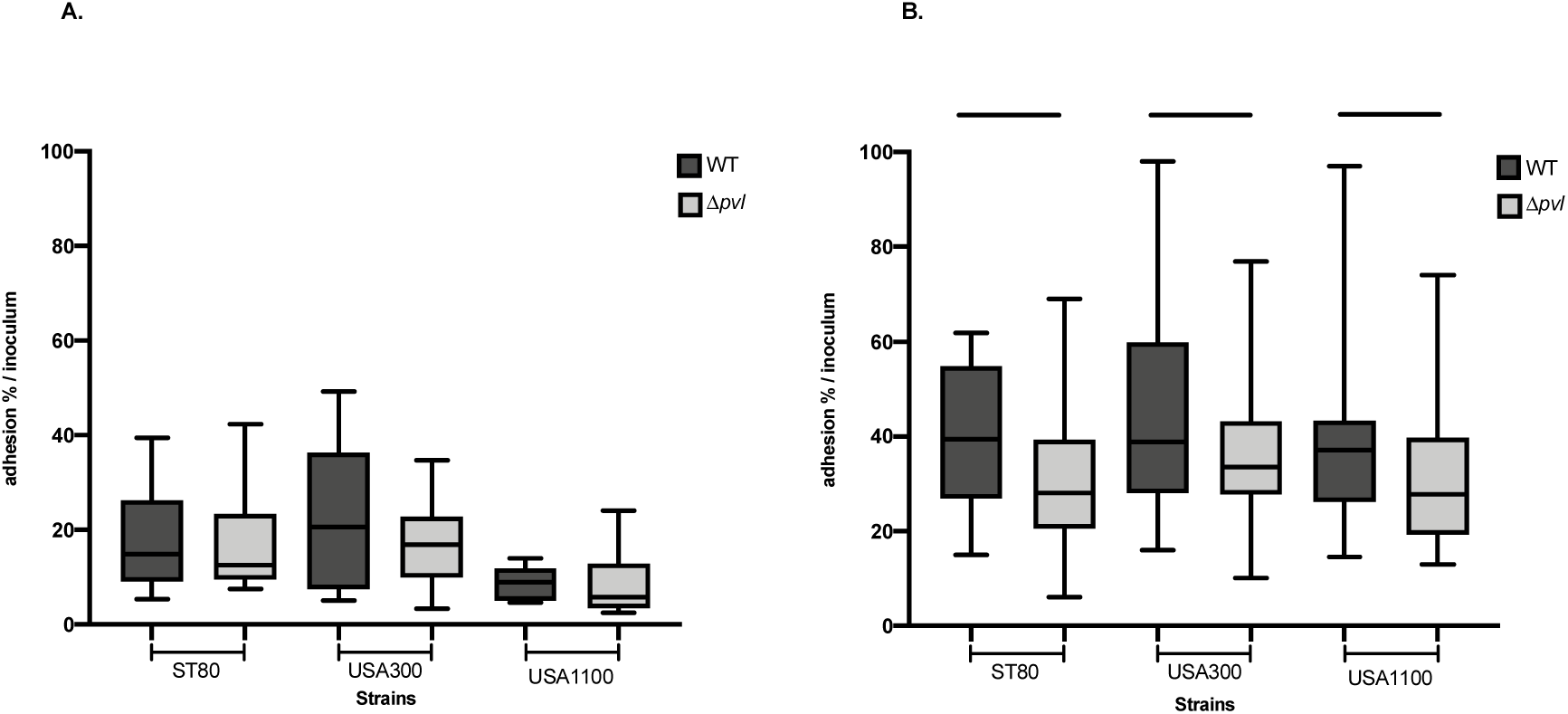
Adhesion of *S. aureus* to intestinal epithelial cells. Three lineages (USA300, USA1100 and ST80) and their isogenic Δ*pvl* derivatives, were tested in a model of bacterial adhesion to intestinal epithelial cells HT29 (panel A) and to mucus-producing intestinal epithelial cells HT29 MTX (panel B). Percentages of adhered bacteria were calculated after 2 hours of infection using as reference the infection inoculum. The horizontal lines within each group represent the mean value ± standard deviations of at least 8 independent experiments per strain. *, *P* < .05; ***, *P* < .001

In multivariate mixed-effect regression, *pvl* presence was significantly associated with an increased adhesion independent of *S. aureus* genetic background or the presence of mucus (Table 3). Including the interaction of *pvl* and mucus presence in the model weakly improved model fit (Akaike information criterion 606.86 compared to 607.45 without interaction) and identified a positive interaction with a magnitude comparable with that of *pvl* in the base model (Table 3), although with a wider uncertainty margin. These results suggest that the positive effect of *pvl* on the adhesion of *S. aureus* to epithelial cells is potentiated by the presence of mucus.

**Table 3.** Mixed-effect multivariate modelling of *S. aureus* adhesion to epithelial cells, with or without interaction between the presence of *pvl* and mucus

| Factor | Odds-ratio of adhesion (95% CI) |  |
| --- | --- | --- |
|  | Base model | Model with *pvl*-mucus interaction term |
| *pvl* presence | 1.49 (1.22 — 1.82) | 1.15 (0.79 — 1.67) |
| Mucus presence | 3.42 (2.65 — 4.42) | 2.86 (2.04 — 4.00) |
| *pvl* and mucus presence | - | 1.44 (0.92 — 2.23) |
| USA1100 (reference) | - | - |
| USA300 | 1.47 (1.15 — 1.90) | 1.47 (1.15 — 1.90) |
| ST80 | 1.07 (0.83 — 1.38) | 1.07 (0.83 — 1.38) |

## Discussion

In the present study we showed that PVL contributes to the colonisation of mouse gut by three CA-MRSA lineages; namely USA300, USA1100, and ST80. We also showed that bacteria were present at the surface of the intestinal mucosa and within the intestinal crypts. Axenic mice were chosen to increase the sensitivity of the colonisation model but do not entirely reflect the reality of bacterial interactions in the gut as its lack normal microbiota. However, the more commonly used antibiotics-treated animal models have also potential draw backs as they shift the microbial population, thus impacting the phenotype studied.^35^ In addition, if such a model had been used, the microbiota could have evolved during the six-week follow-up of the study in an uncontrolled manner. As we compared isogenic strains belonging to three lineages, we believe the results obtained are consistent and likely reflect a relevant interaction of *S. aureus* with the mucosal surface mediated by PVL. The hypothesis that PVL could contribute to other functions beyond pore formation was raised previously, following the observation that the signal peptide of the PVL LukS component enhances bacterial adhesion to matrix protein and to heparan sulfates which are highly abundant at the mucosal surface.^36^ Whatever the precise mechanism underlying bacterial adhesion to the mucus, adhesion to intestinal epithelial cells by the three tested strains appeared to be PVL-dependent and mucus-dependent, as a difference in adhesion between WT and isogenic *pvl* knockout was only observed on mucus-producing cells. The possibility that USA300, USA1100, and ST80 wild type outcompeted their Δ*pvl* derivatives in the mouse model by direct bacterial interference was ruled out by the *in vitro* competition experiment which showed no fitness disequilibrium. Moreover, WT and isogenic *pvl* knockout strains isolated after 42 days from the mice gut and further challenged in the *in vitro* competition assay also did not reveal an altered fitness of the Δ*pvl* derivative. This allowed to ensure that the results of the challenge in the mouse model were not a consequence of deleterious mutations acquired, during the 42-day experiment, by the Δ*pvl* strain. Taken together these observations support the hypothesis that PVL contributes to CA-MRSA gut colonisation by enhancing interactions with the mucus. The intestinal mucosal barrier plays a key role in intestinal homeostasis notably by preventing translocation of pathogenic bacteria. Intestinal bacterial overgrowth and dysbiosis are associated with alteration of this barrier.^37^ Here, because of the use of an axenic mouse model of gut colonisation, we could not demonstrate whether *S. aureus* colonisation of the mucosal layer had any deleterious consequences on gut microbiota and/or on intestinal permeability. However, the fact that intracellular bacteria were not found on histological sections of the caecum, nor in the intestinal cellular model (not shown), suggests that *S. aureus* does not demonstrate a potential for invading these cells and rather behave as a bystander in the mucus, resulting in a fully commensal relationship with the intestinal mucosal barrier. Although this working model needs to be deciphered further, we feel that given the complexity and considerable duration of the mice experiments, it is important to disclose these results which shed new light on a possible contributing factor to the success of CA-MRSA; that is the enhancement of CA-MRSA gastro-intestinal colonisation possibly mediated by a moonlighting function of PVL.

## Acknowledgment

The authors thank Anabelle Bouchardon (CIQLE) for technical help in microscopy imaging and Sandra Hoys for technical help in mice experiments. We are thankful to Nicolas Barnich, Paul Verhoeven and Karen Moreau for valuable scientific advices and to Véréna Landel from the Hospices Civils de Lyon for his precious guidance in the writing of the manuscript and his expertise in English spelling.

## Transparency declarations

None to declare

